# POTATO: An automated pipeline for batch analysis of optical tweezers data

**DOI:** 10.1101/2021.11.11.468103

**Authors:** Stefan Buck, Lukas Pekarek, Neva Caliskan

## Abstract

Optical tweezers is a single-molecule technique that allows probing of intra- and intermolecular interactions that govern complex biological processes involving molecular motors, protein–nucleic acid interactions and protein/RNA folding. Recent developments in instrumentation eased and accelerated optical tweezers data acquisition, but analysis of the data remains challenging. Here, to enable high-throughput data analysis, we developed an automated python-based analysis pipeline called POTATO (<u>P</u>ractical <u>O</u>ptical <u>T</u>weezers <u>A</u>nalysis <u>TO</u>ol). POTATO automatically processes the high-frequency raw data generated by force-ramp experiments and identifies (un)folding events using predefined parameters. After segmentation of the force-distance trajectories at the identified (un)folding events, sections of the curve can be fitted independently to worm-like chain and freely-jointed chain models, and the work applied on the molecule can be calculated by numerical integration. Furthermore, the tool allows plotting of constant force data and fitting of the Gaussian distance distribution over time. All these features are wrapped in a user-friendly graphical interface (https://github.com/REMI-HIRI/POTATO), which allows researchers without programming knowledge to perform sophisticated data analysis.

**SIGNIFICANCE:** Studying (un)folding of biopolymer structures with optical tweezers under different conditions generates very large datasets for statistical data analysis. Recent technical improvements accelerated data acquisition by coupling modern instruments with microfluidic systems, at the same time creating the need for a high-throughput, and unbiased data analysis. We developed Practical Optical Tweezers Analysis TOol (POTATO); an open-source python-based tool that can process data gathered by any OT force-ramp experiment in an automated fashion. POTATO is principally designed for data preprocessing, identification of (un)folding events and the fitting of the force-distance curves. In addition, all parameters for preprocessing, statistical analysis and fitting of the curves can be adapted to suit the dataset under analysis in an easy-to-use graphical user interface.

## INTRODUCTION

Arthur Ashkin received the Nobel Prize in 2018 for his research on trapping dielectric particles with laser light in optical tweezers (OT) (1). Optical tweezers enable probing of structural dynamics of individual molecules by monitoring internal forces and short-lived intermediate states in real-time (2–5). This technique has been widely used to study structures of nucleic acids and dynamics of RNA/protein folding (6–10). In addition, OT can also be used to probe the molecular interactions between small molecules, proteins, and nucleic acids (11–13). Recently, the combination of optical tweezers with confocal microscopy enabled simultaneous measurements of force and fluorescence that provided unprecedented insights into molecular mechanisms such as timing and order of events during transcription or translation (12,14–16). Basically, in a typical OT experiment, a biopolymer, such as a protein, DNA, or RNA molecule, is tethered between two dielectric beads via labeled handles. The beads are then trapped by focused laser beams, the so-called optical traps. Following this several modes of operation are possible. In force-ramp mode the beads are precisely displaced in a monotonous manner, which applies increasing forces onto the biopolymer (**Fig. 1A**). Since trapped beads behave as if they were attached to mechanical springs, the applied force can be calculated from the measured end to end displacement of the beads out of the trap focus according to Hooke’s law (**Fig. 1B**) (17). This mode is commonly used to determine the elastic properties of the molecule and/or to determine the rupture forces at which transitions in folding and unfolding occur. On the other hand, a constant-force operation mode allows tracking the molecule of interest in real time as it transitions between different conformational states, yielding kinetic parameters of folding-unfolding of molecules or progressive movements of molecular motors (5). Accordingly, optical tweezers experiments also allow precise calculation of the work done on the system of interest (18,19).

**FIGURE 1:**
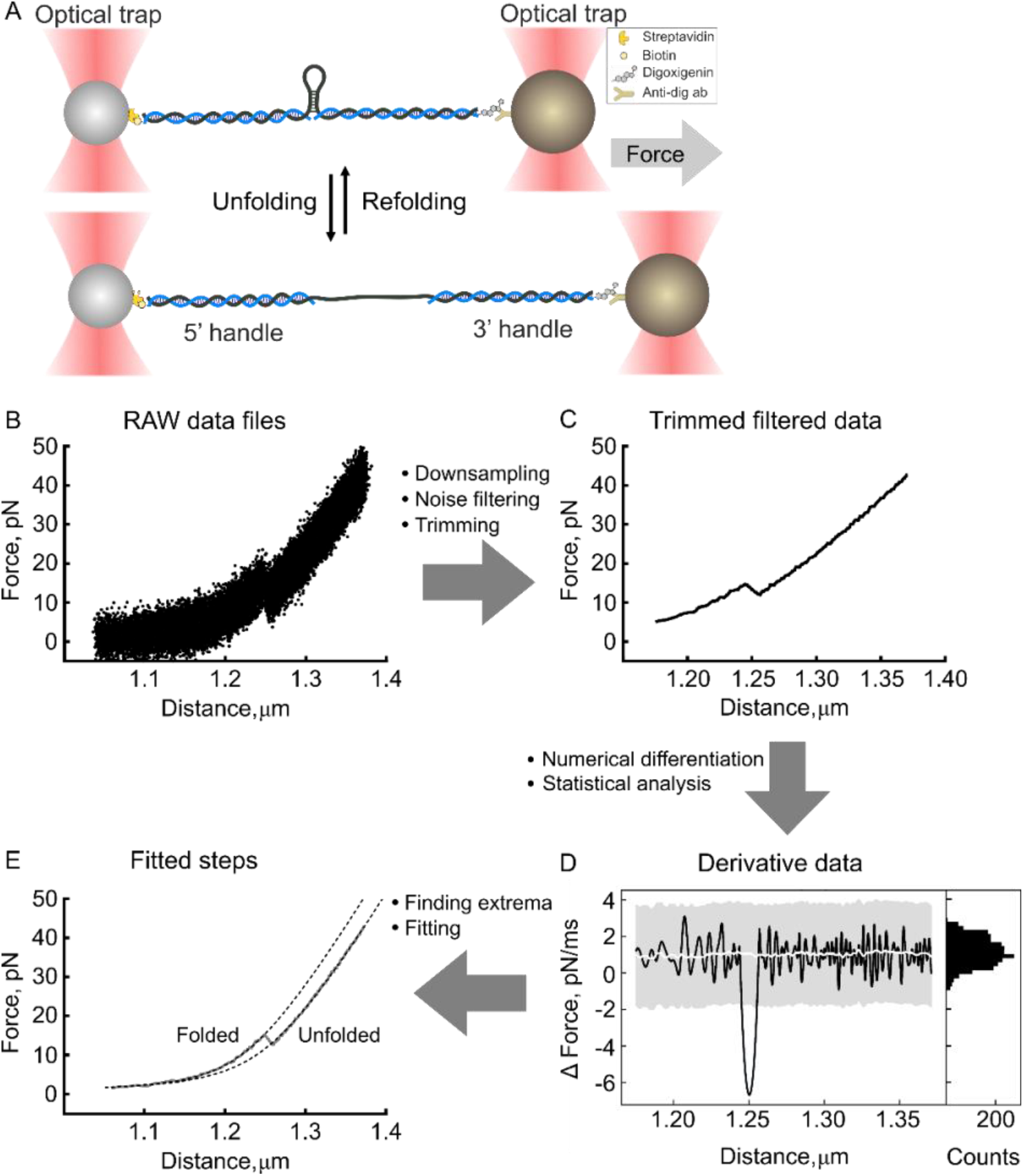
Schematic of the pipeline. **(A)** Diagram illustrating the optical tweezers experiments. RNA is hybridized to ssDNA handles and immobilized on beads. These are used to exert a pulling force on the RNA with a focused laser beam. In force-ramp operation mode, the force is gradually increased until the structure in the middle is unfolded (bottom). *RAW data files* **(B)** are down sampled, the noise is filtered using a Butterworth signal filter, and the data are trimmed at a minimum force threshold to yield the *Trimmed filtered data* **(C)**. Then the time derivative is calculated numerically to yield the *Derivative data* **(D)**; histogram of the derivative value distribution (right) shows two populations - normal-like distribution represents the experimental noise, while the other population of outliers represents the (un)folding steps. The derivative data are then statistically analyzed – the standard deviation and moving median are calculated. Peaks in derivative data that exceed median (white line) ± z-score (grey region) are classified as (un)folding events. The beginning and end of each event are indicated. The coordinates of the events are then used to define the region for fitting, yielding the *Fitted steps* **(E)**. Finally, the output data files are exported according to the selected settings.

Previously, OT instruments were self-built by researchers and thus application required substantial physics and engineering background. Furthermore, such experiments were highly time demanding and labor intensive because a large amount of data need to be collected for a quantitative analysis. Recently, commercial instruments became available on the market. Another breakthrough in the field was the integration of OT instruments with microfluidic systems, which accelerated both experimental setup and data acquisition (14,15). Nowadays, high-frequency data acquisition allows the generation of large data sets in a relatively short time. Subsequent data analysis, however, still requires custom written scripts to perform data preprocessing, identification of (un)folding events or different folding states, mathematical modeling, and statistical analysis. Although device manufacturers provide basic scripts for the analysis of experimental data, data processing still requires bioinformatics and statistics skills and thus remains being a major bottleneck.

Here, we present an automated python-based pipeline for optical tweezers force-ramp and constant-force data analysis. Using statistical analysis of the time-derivative of force and distance data, (un)folding steps are automatically identified, and values such as (un)folding force and step length are derived. These values are then directly used for fitting of force-distance (FD) curves. Additionally, we provide a basic constant-force analysis function. We have also integrated an easy-to-use graphical user interface (GUI), allowing users to change the analysis parameters to suit their needs (**Table S1**). Importantly, this workflow ensures reproducibility and eliminates inconsistencies of manual analysis (20). Since the pipeline allows automated processing of multiple raw data files, analysis time is substantially reduced. Finally, applicability of this tool is demonstrated on an artificially generated dataset, which covers a broad range of possible parameter combinations for force-ramp data, and also real experimental data (21). Our results indicate that ‘POTATO’ exhibits a robust performance in identifying (un)folding events with high accuracy, precision and recall.

## MATERIALS AND METHODS

### Algorithm implementation

The algorithm is written in python 3. We designed a graphical user interface and wrapped the code into a windows stand-alone executable with *pyinstaller* to open this tool to a broader audience without a bioinformatics background. The code can be found on GitHub (https://github.com/REMI-HIRI/POTATO) and the architecture of the python files and GUI is further explained in the Supporting Material.

### Artificial data generation

Artificial force spectroscopy data were generated using a custom-written python script (Supporting Material). The fully folded part of FD curves was modeled using an equation for extensible worm-like chain (WLC) models (**Eq. 4**). The partially unfolded region was modeled using a combination of WLC and freely-jointed chain (FJC) models (**Eq. 5 and 6**). For a more detailed description, see the supplementary information.

### Optical trapping system

Optical tweezers experiments were performed using a C-Trap® instrument (LUMICKS, Amsterdam/NL). This device offers two laser traps combined with a 5-channel laminar-flow microfluidics system and a confocal microscope. Experiments were conducted as described in (21,22).

### RESULTS AND DISCUSSION

### Data preprocessing

Raw data (**Fig. 1B**) from various input file formats (**Supporting Material**) can be loaded and preprocessed before marking unfolding events. Initially, the data are down sampled. This is especially important when data are collected at high frequencies, because down-sampling accelerates the analysis and saves storage space. Then, a low pass Butterworth filter is used to reduce the noise of the signal (**Eq. 1**) (23). This filter allows efficient noise removal while keeping the actual (un)folding events intact and is therefore commonly used (**Fig. 1C**). The algorithm then trims the data at a minimum force threshold set by the user (**Table S1**).

(1) Butterworth filter:

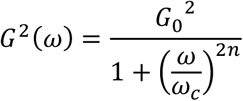

*G* is gain, *ꙍ* is frequency, *ꙍ*_*c*_ is cut-off frequency, and *n* is filter degree.

### Force-ramp data analysis

#### Statistical analysis

In force-ramp trajectories, an (un)folding event is characterized by a simultaneous drop in force and a quick increase in distance as a secondary structure of the polymer undergoes a sudden transition from the folded to the unfolded state (**Fig. 1C**). This (un)folding event can be identified as a local maximum in the derivative of the distance and a local minimum in the derivative of the force (**Eq. 2**). When plotted, the numerical derivative data of both distance and force show two populations of values. The first is a normal-like distribution representing the measurement noise, while outliers from the normal distribution represent the second population – the actual (un)folding events. To distinguish real (un)folding events from background noise, we calculate the moving median and the standard deviation (SD). These are then used to separate the normally distributed data from the extreme values outside a given z-score (i.e. number of standard deviations, 3 by default) (**Fig. 1D**). This should include 99.73% of the normally distributed data points. As the initially calculated SD is affected by the outliers, a second SD is calculated from the data points inside the threshold, and the data are sorted again. The cycle is repeated until the difference between initial and secondary SD is < x (with x-default = 5%). After the force- and distance derivatives are sorted, our algorithm finds the local extrema of the derivatives, representing the saddle points of the (un)folding events in the FD curve. Then, it finds the adjacent crossing points of the derivative with the moving median, representing the start or end of the corresponding unfolding events.

(2) Numerical approximation of the derivatives:

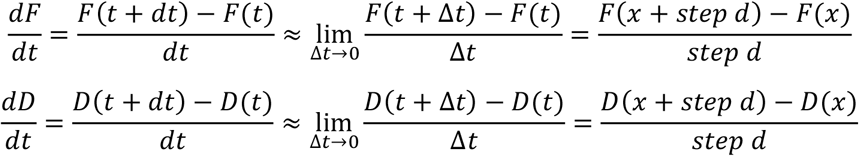

F is force, D is distance, *t* is time, *x* is position, and step *d* is a change in position.

#### Data fitting

After (un)folding steps are identified, this information can be used for data fitting. For the characterization of the mechanical properties of the (bio)polymer under tension, the worm-like chain (WLC) model is commonly used (**Eq. 3**). Briefly, the FD curve is split into multiple parts. The fully folded part (until the first detectable unfolding step) is fitted with a worm-like chain model (WLC) (24) to determine the persistence length (dsL_P_) of the tethered molecule, while the contour length (dsL_C_) is fixed. The other (partially) unfolded parts of the FD curve are then fitted by a combined model comprising WLC (describing the folded double-stranded handles) and freely jointed chain (FJC) (**Eq. 4, 5**), or another worm-like chain (WLC) model (representing the unfolded single-stranded parts) (**Eq. 6**) (**Fig. 1E**) (24,25). To mathematically fit the models, we use the *pylake* (LUMICKS) python package that contains the fitting functions with commonly used variants of the model equations.

(3) Worm-like chain model (WLC):

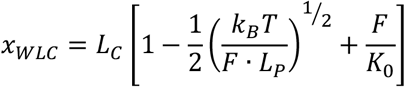

*X* is an extension, *L*_*C*_ is contour length, *F* is force, *L*_*P*_ is persistence length, *k*_*B*_ is Boltzmann constant, *T* is thermodynamic temperature, and *K*_*0*_ is stretch modulus.

(4) Freely jointed chain (FJC):

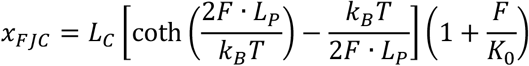

(5) WLC + FJC:

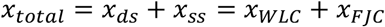

(6) WLC + WLC:

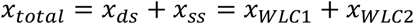

#### Work calculation

Unfolding and refolding force-distance trajectories also yield crucial information on the thermodynamic properties of the molecule under study. Accordingly, the work applied by the optical tweezers instrument onto the system can be calculated from the area under the FD curve (AUC), here using composite Simpson’s rule (**Eq. 7**). The calculated area gives the work applied to the whole construct, including the handles. Thus, in order to determine the amount of work applied to the structure of interest, the work applied to the handles, represented by the AUC until the end of the step of the combined model, is subtracted (**Eq. 8**). It shall be noted that the work derived from these calculations equals Gibbs free energy of the studied structure when the system is close to the thermodynamic equilibrium. However, typically the (un)folding trajectories may not coincide indicating that the molecule is out of equilibrium. Despite that even when the system is not in equilibrium, Gibbs free energy can be extracted from the work values (5,18,19,26,27).

(7) Numerical integration using composite Simpson’s rule:

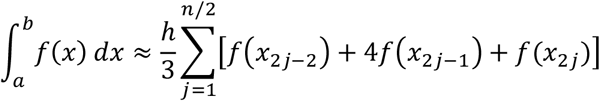

where *x*_*j*_ = *a* + *jh for j*=*0, 1*, …, *n-1* with *h*=*(b-a)*/*n*; *x*_*0*_ = *a* and *x*_*n*_ = *b*.

(8) Non-equilibrium work calculation:

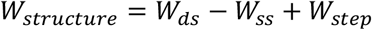

W_structure_ is work needed to unfold the structure of interest. W_ds_ is numerical integration of the fully folded model, W_ss_ is numerical integration of the unfolded model, and W_step_ is numerical integration of the step region between the two models.

### Constant force data analysis

In addition to force ramp experiments, the algorithm we provide can also analyze constant force data (**Fig. S1** in the Supporting Material). In this way, the behavior of the structure around the (un)folding force can be investigated, and the equilibrium force at which the chance of the structure to be folded or unfolded are equal can be derived.

The constant force analysis accepts the same input formats as the force-ramp batch analysis, and data preprocessing is performed similarly by down sampling and filtering, without trimming. First, it is necessary to display the constant force data in order to optimize the preprocessing parameters and the plot’s axis (**Fig. S1B)**. At this step, two plots are generated for visualization. In the first plot, distance is plotted against time. Here, the difference in distance corresponds to the change in the contour length of the tethered molecule. The second plot is a histogram of the distance distribution (**Fig. S1C)**. From this histogram, the number of different folding states can be deduced. Afterward, the histogram can be fitted with multiple Gaussian functions. According to the position distribution histograms, the user can interactively provide initial estimates for various parameters including the number, localization, width (standard deviation, z-score), and amplitude of the fits. After the optimization, the optimized parameters are exported together with the percentage of each folding state as a table in csv (comma separated values).

### Artificial data sets to test the limits of detection

To test the limits of (un)folding events detectable by the POTATO pipeline, an artificial dataset was generated (Supporting Material). In this data set, some curves can show a negative step-length that would not be observed in real unfolding events. We considered these steps as non-identifiable and used them as negative controls. The phenomenon of negative steps can mainly be observed for small contour length changes (ΔL_C_) between the models, combined with high force drop (ΔF) values. To test the performance of the algorithm, we defined identifiable steps as events with a drop in force and a simultaneous increase in distance (Supporting Material). To evaluate if a specific parameter combination results in an identifiable curve, **Eq. 9** with x = 0 was solved for all sets of parameters. Each time two parameters were fixed, and the third parameter was optimized.

(9) Minimal step calculation:

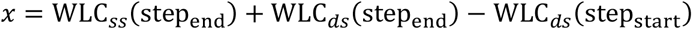

Where WLC corresponds to expression from **eq. 3**, “ss” refers to the model corresponding to single-strand values, while “ds” describes the double-stranded region.

A hyperplane showing the interface of theoretically identifiable and non-identifiable steps was generated from these optimized values (**Fig. 2A**). This allowed us to classify the generated dataset based on a combination of parameters: One with curves where POTATO is expected to find an unfolding step (x > 0) and the other one where POTATO should not identify the steps (x ≤ 0). After analyzing the artificial dataset (comprising 2520 curves) with different z-scores, the expected results, based on the input parameters when the data were generated, were compared to the steps identified by POTATO. For the default z-score of 3, the expected parameters were then plotted into the 3D plot and colored based on the identification by POTATO (**Fig. 2A**). For an unfolding force of 25 pN, the ΔF and ΔL_C_ values are shown in a 2D plot, making it easier to identify and compare single unfolding events analyzed with different z-scores. It can be seen that all identified steps at this specific unfolding force are above the theoretical threshold and that more unfolding events are identified at z-score 2.5 than at z-score 3 (**Fig. 2B**). Accordingly, the effect of the z-score on the derivative of force (**Fig. 2C)** and distance (**Fig. 2D**) can be investigated for an individual force-distance trajectory. In the representative trajectory, the local maximum in the derivatives of distance is above the z-score threshold for both cases. In the derivative of force, the local minimum at the same position is only detected for the lower z-score **(Fig. 2C-D)**.

**FIGURE 2:**
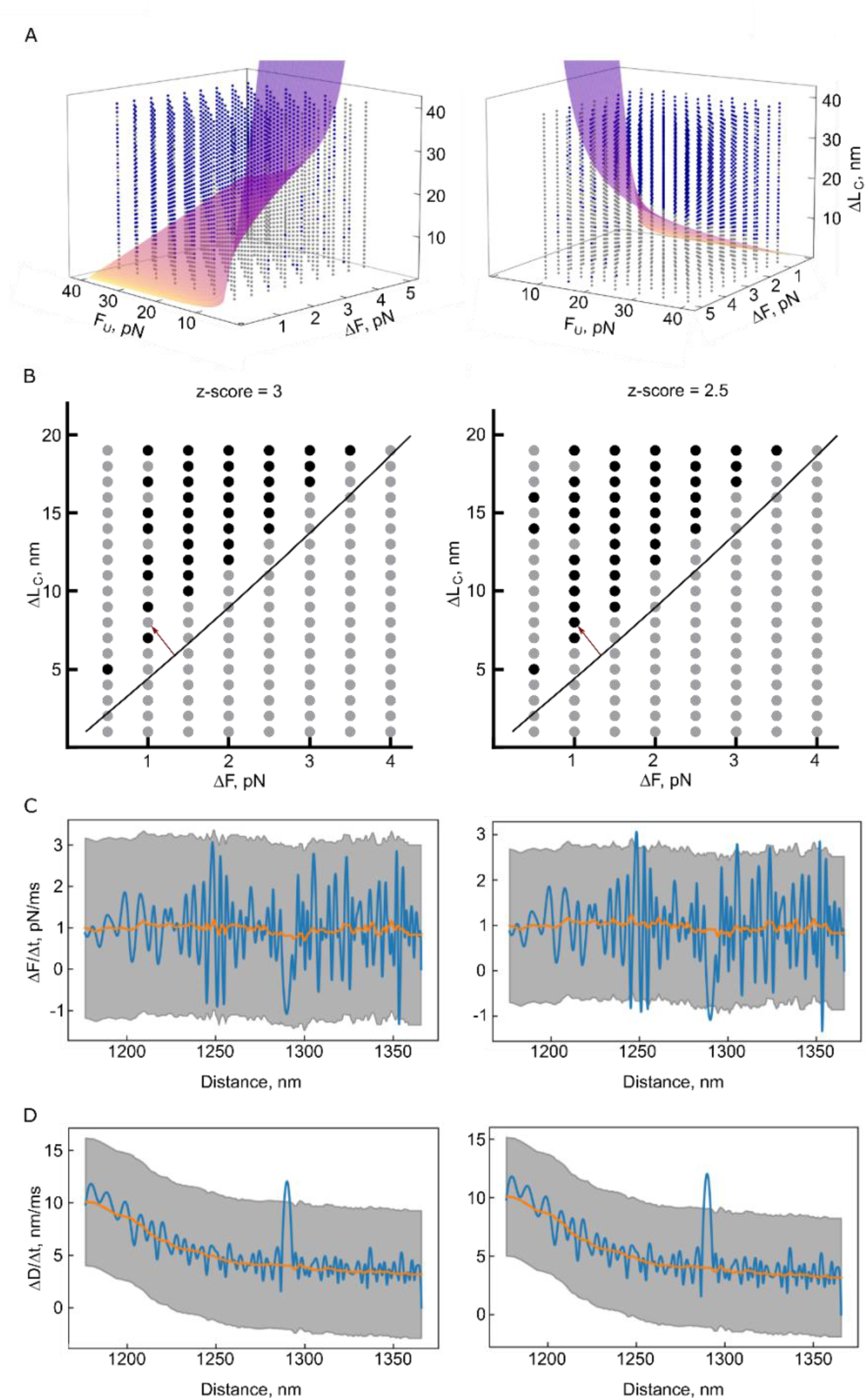
Testing the limits of POTATO. For each combination of the parameters unfolding force (F_U_), force drop (ΔF), and contour length change (L_C_), two parameters were fixed, and the third one was optimized so that the **eq. 9** (Supporting Material) evaluates to zero. **(A)** A hyperplane was generated from the optimized values that separate the resolvable space above the hyperplane (parameter combinations that result in identifiable steps) from the unresolvable space below the hyperplane (parameter combinations that result in unidentifiable steps). Each analyzed curve is plotted in blue if its step was identified by POTATO or in grey if it was not recognized. **(B)** Slices of the 3D plot at F_U_ = 25 pN were analyzed with different z-scores. The black line corresponds to the theoretical limit of resolvable/unresolvable parameter combinations. The black dots represent curves with identified steps, whereas the grey dots represent curves where POTATO could not identify the step. The derivatives of force **(C)** and distance **(D)** of the curve that is marked with a red arrow in **(B)** are displayed at different z-scores.

Next, we calculated performance measures such as accuracy, precision, sensitivity, specificity, and F1-score to validate the performance of POTATO. For a z-score of 3.2, a precision score of 0.974 indicates that most of the positive classified steps were actual steps, and even for a z-score of 2.5, the precision was still above 0.944 (**Table S2**). As expected, higher precision comes with the trade-off to miss certain positive events (recall 0.870 - 0.939), and the optimal z-score has to be chosen depending on the application. For smaller unfolding events that are difficult to detect, lower z-scores should be employed, as for distinct unfolding events the z-score can be set to higher values. This way number of false-positive events detected can be minimized. Since the present dataset was generated using artificial parameter combinations, those might not be found in actual OT measurements. Therefore, it is important to keep in mind that we were exploring the limits of the tool by using these strict parameter constraints. Performance measures would also vary depending on where a specific dataset is located in the parameter space, and which z-scores were employed.

Furthermore, we investigated how accurately POTATO estimates step parameters (F_U_, ΔL_C_, ΔF). For that, we compared the expected and measured values of these parameters for all curves analyzed (**Fig. 3**). We then calculated the linear regression of the true positive values to estimate possible biases of POTATO estimated F_U_ and ΔL_C_ values. Our analysis shows that in the case of F_U_ (**Fig. 3A**), the values determined by POTATO are in perfect agreement with the expected values (slope of the linear regression = 0.9912). For ΔL_C_ (**Fig. 3B**), the comparison shows a broader distribution of the measured values with an overall trend suggesting a minor overestimation (slope of the linear regression = 1.0282) of around 3%. Because we observed several true positives highly under- or overestimating the ΔL_C_, we further explored these events, which are likely caused by wide ranges of the fitting constraints. This phenomenon can be strongly minimized by adapting the fitting constraints like the stiffness or persistence length boundaries. Lastly, in the case of ΔF (**Fig. 3C**), the trend shows a slight underestimation of the measured values (slope of the linear regression = 0.8517), resulting in a bias of roughly 15%. Taken together, our performance measures analysis suggests that the presented tool successfully identifies most (un)folding events correctly with only few false classifications (false positives/false negatives). Accordingly, in most of the cases, performance measures were above 0.9 (**Table S2**). Moreover, we show that POTATO can precisely estimate the parameter values describing the (un)folding events (F_U_, ΔL_C_, ΔF, **Fig. 2**). Overall, the performance measures and the accuracy of the estimates show that POTATO represents a reliable tool for optical tweezer data analysis.

**FIGURE 3:**
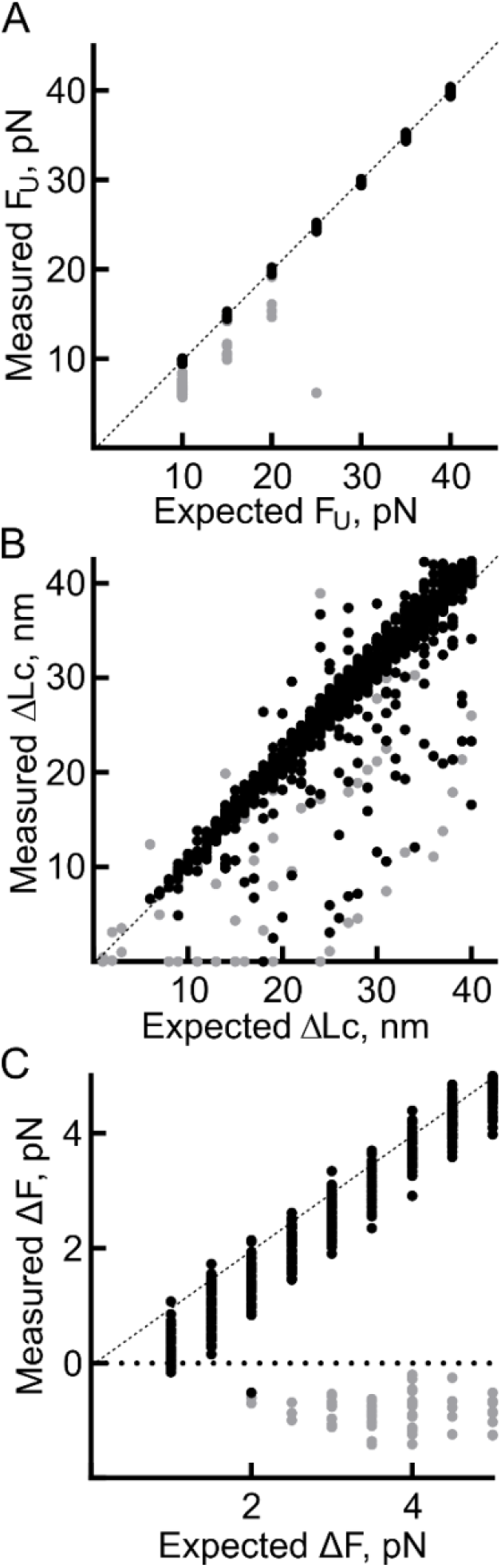
Evaluation of the performance of POTATO. The parameters used for the generation of the dataset compared to the parameters identified by POTATO are plotted against each other. All three parameters used for the data generation are evaluated with a z-score of 3. The values of the true positive steps (black) and the values of the false-positive steps (grey) are visualized for **(A)** the unfolding Force (F_U_), **(B)** the contour length change (ΔL_C_), and **(C)** the force drop (ΔF). A dashed line represents the theoretical perfect correlation between measured and expected values.

### Applicability of POTATO on real experimental data

Next, we employed POTATO to test its performance on real experimental data generated from force-distance measurements of the programmed ribosomal frameshifting (PRF) element of the Encephalomyocarditis virus (EMCV) (21,28). We compared the POTATO results with manually annotated steps of a subset of our dataset. The results obtained with manual step identification and data fitting were in good agreement with the automated analysis using the pipeline (**Fig. S2** in the Supporting Material). Harnessing POTATO in the data processing allowed us to speed up the analysis significantly compared to previous manual analysis. Furthermore, we saw that POTATO is not only suitable for curves with a single (un)folding event like in the artificial dataset, but we successfully fit force distance curves with as many as five unfolding steps and we were able to identify even short-lived intermediate states of the unfolding process (**Fig. S2B** and **C**). In addition to the contour length change obtained by curve fitting, also the Gibb’s free energy is an important variable to conclude on the nature of the (un)folded structure as the Gibb’s free energy is dependent on the base pairing of the RNA. With this experimental dataset we were able to use the work calculated by the analysis pipeline, to estimate the Gibb’s free energy of the structures and thereby distinguish between different secondary structures (18,27). The findings on these data will be published in a separate paper, which is accepted for publication (21). In conclusion, we showed that the pipeline output matches with manual data analysis on real-experiment data and that the tool we present performs analysis of FD trajectories with multiple steps or even short-live intermediates. Therefore, POTATO represents a versatile tool for high-throughput OT data analysis for many upcoming studies. **Limitations**

Processing automation comes with trade-offs (29,30). First, the statistical analysis applied in the pipeline might be prone to false-positive event discoveries due to external causes, such as vibration that might induce step-like events in the force-distance profile of gathered data. We split the force-distance data and analyze the derivatives of force and distance separately to minimize this effect. Only the events found by both approaches are considered real (un)folding events. Therefore, the robustness of the analysis is increased. Second, the pipeline output strongly depends on parameters and threshold values that are applied throughout the analysis. The default values were set empirically to suit our needs. Therefore, it might require optimization to fit specific needs and reach an analysis output consistent with the manual data analysis. User input is still required despite the user-friendly GUI environment, and an understanding of the analysis workflow is necessary to adjust the parameters rationally. Lastly, the current algorithm does not annotate the repeated folding and unfolding of a structure during force-ramp measurements and identifies this oscillation as independent steps. Nevertheless, this mainly occurs at slow loading rates and does not affect the contour length estimates.

## SUMMARY

Here we present a publicly available pipeline for batch analysis of optical tweezers data. Our pipeline allows OT raw or preprocessed data processing from force-ramp or equilibrium measurements (constant force/position). These are widely employed experimental approaches in the OT field, applied to nucleic acid structure probing, protein folding, RNA-protein interactions, or even to analyze events as complex as translation. Here, by wrapping our algorithm in a standalone application and designing an intuitive graphical user interface, we aim to open the data analysis to a broader audience without the need for a bioinformatics background. The user can adjust all parameters directly in the GUI without diving into the code to tailor the pipeline to their exact needs. With the parameters optimized for the here presented datasets, POTATO showed high precision and accuracy in the identification of (un)folding events. Moreover, compared to manual data analysis, the pipeline is faster and, most importantly, consistent throughout the analysis, thus yielding reproducible results.

## SUPPORTING MATERIAL

Supporting Material can be accessed in the GitHub repository (https://github.com/REMI-HIRI/POTATO).

## AUTHOR CONTRIBUTIONS

NC, LP, and SB designed the pipeline. LP and SB wrote the python scripts. LP generated the artificial data. SB analyzed the artificial data. LP and SB performed the optical tweezers experiments. LP analyzed experimental data. LP and SB prepared the figures with the input from NC. NC, LP, and SB wrote the manuscript.

## ACKNOWLEDGMENTS

We thank Vojtech Vrba for helpful python discussions. We thank Dr. Anke Sparmann for critically reviewing the manuscript. The work in our laboratory is supported by the Helmholtz Association and grants from the European Research Council (ERC) Grant Nr. 948636.

## SUPPORTING CITATIONS

**References (31–35)** appear in the Supporting Material.

