## Supplementary information for "POTATO: An automated pipeline for batch analysis of optical tweezers data"

### Script structure

The script is written in Python 3 and split into multiple parts for clarity. The first part, "POTATO_GUI", defines the GUI with all necessary functions and input variables. When the GUI is started, the default values of the input variables are loaded from the "POTATO_config" file. The GUI was created and structured using the standard Tkinter python package. A parallel subprocess initiates from this main process when a folder is selected for force ramp analysis to perform computationally demanding data-processing. This way the GUI remains responsive during computation. All the functions used for data preprocessing and step recognition are defined in the "POTATO_preprocessing" and the "POTATO_find_steps" files respectively. The functions used for curve fitting are defined in another file, "POTATO_fitting". For computation, we mainly use matplotlib and NumPy packages, as well as the lumicks.pylake package for fitting (**Table S3**). The subprocess is a daemon process spawned by the main process and therefore stops as soon as the GUI terminates the main process. The last part, "POTATO_constantF", is executed by the main thread as it only analyzes one constant force file at a time. The results are exported in different CSV files or as PNG images.

### Graphical user interface

We designed a graphical user interface (GUI) that allows users to easily adjust the analysis steps and parameters according to their needs and select between three different input data formats. This enables the GUI to load data from every OT instrument. The GUI is separated into multiple tabs, resulting in easy and intuitive navigation without overloading the individual windows. The "POTATO_config" file, included in the POTATO repository, contains the default parameters, which are loaded into the GUI. The most commonly changed parameters can be found in the first tab, “Folder Analysis”, so a basic analysis can be performed right away (press enter to confirm changed parameters). Alternatively, before each analysis, all parameters can be adjusted in the 'Advanced Settings' tab to suit the data set. In addition, we implemented the possibility to selectively export results. Each analysis creates a new folder with a timestamp directly in the analyzed directory. The used parameters are exported as well so that parameters can be optimized later. The second tab, “Show Single File”, provides a control mechanism for data preprocessing. A single file can be loaded, and the unfiltered data are plotted together with the filtered data, which streamlines troubleshooting. Finally, there is a third tab for “Constant Force Analysis”.

### Input data

The presented pipeline accepts three different input data formats. Two of them are based on the default hdf5 output format of Lumicks C-Trap – one is predefined for high-frequency data (using the piezo-tracking function of the instrument), and the second is for low-frequency data (using video recognition). The third data format is a basic CSV file format with force and distance values in the first and second columns. Thus, our pipeline can process force-distance data from virtually any optical tweezers machine. In addition, entire directories containing force-ramp data files can be selected and processed simultaneously.

**Data output**

Depending on individual analysis requirements, different export settings can be selected. The down sampled and filtered data are exported in CSV format (smooth) for each file by default. The identified (un)folding steps by derivatives of force and distance are exported together with the steps identified by both strategies (common steps) into a single CSV file. All identified steps of all curves in the analyzed folder are gathered into a single results file for quantitative analysis. The respective summary figure containing the plot of preprocessed data, trimmed data, and both derivatives with marked steps is exported. The plots of fitted data, together with the fitting parameters and model data, are exported as PNG and CSV files, respectively.

**Artificial data generation**

To test the limits of the algorithm, artificial data with a single step per curve were generated. The fully folded part of force-distance curves was modeled using an equation for extensible WLC models (**Eq. 4**). The partially unfolded region was modeled using a combination of WLC and FJC models (**Eq. 5 and 6**). The force value at which the step occurs, the contour length change between the unfolded and folded region, and the drop in force during the step, are the parameters for data generation. The first parameter was set to occur between 10-40 pN with a 5 pN resolution. The curves were generated with a contour length change from 1-40 nm with a 1 nm resolution and a force drop of 1-5 pN with a 0.5 pN resolution. To mimic the (Gaussian) noise affecting the raw data, we employed the NumPy random normal distribution function (1).

Since the (un)folding step is generally defined as a drop in force (one of the parameters) and a sudden increase in distance (not a parameter), the data generated by this script also contained combinations that did not increase distance. We used these curves showing no increase in the distance as negative controls.

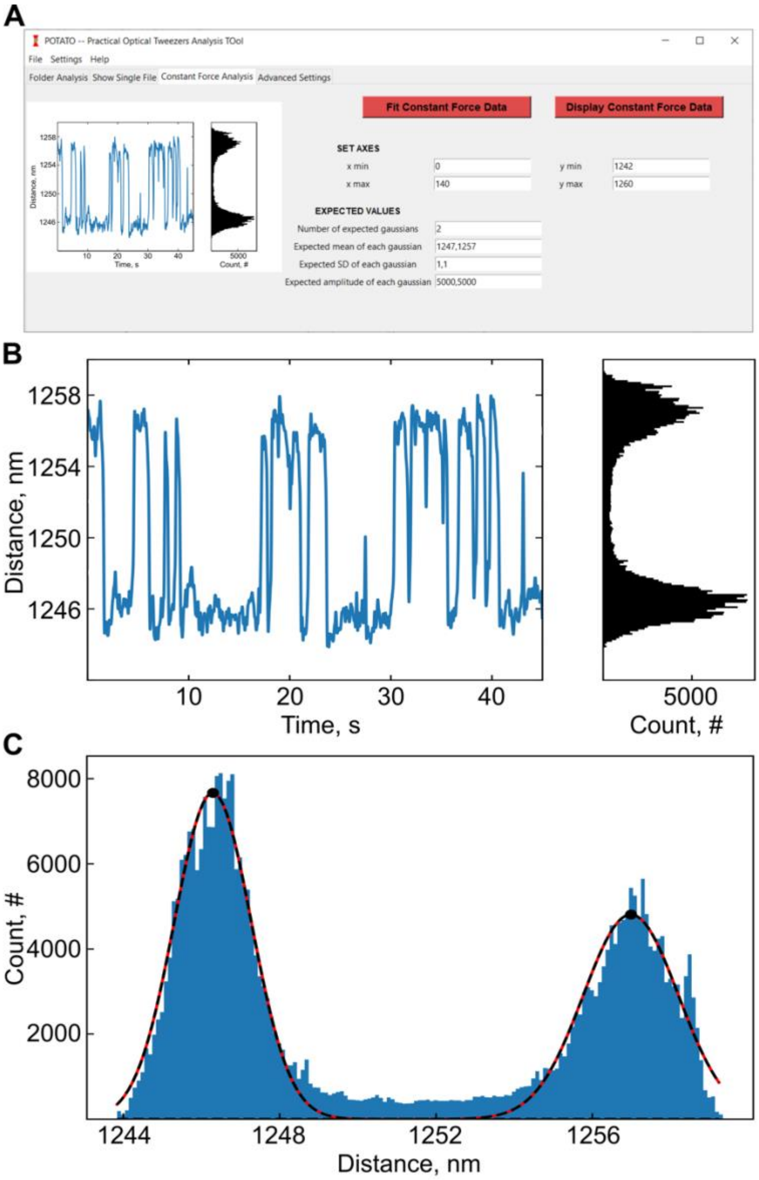


**FIGURE S1: Constant force data analysis in POTATO: (A)** GUI tab containing the constant force analysis features, **(B)** Display constant force data output; (left) distance over time plot, (right) histogram of the distance over time values. **(C)** Fit constant force data output showing the histogram of distance values distributions and the two gaussian functions fitted.

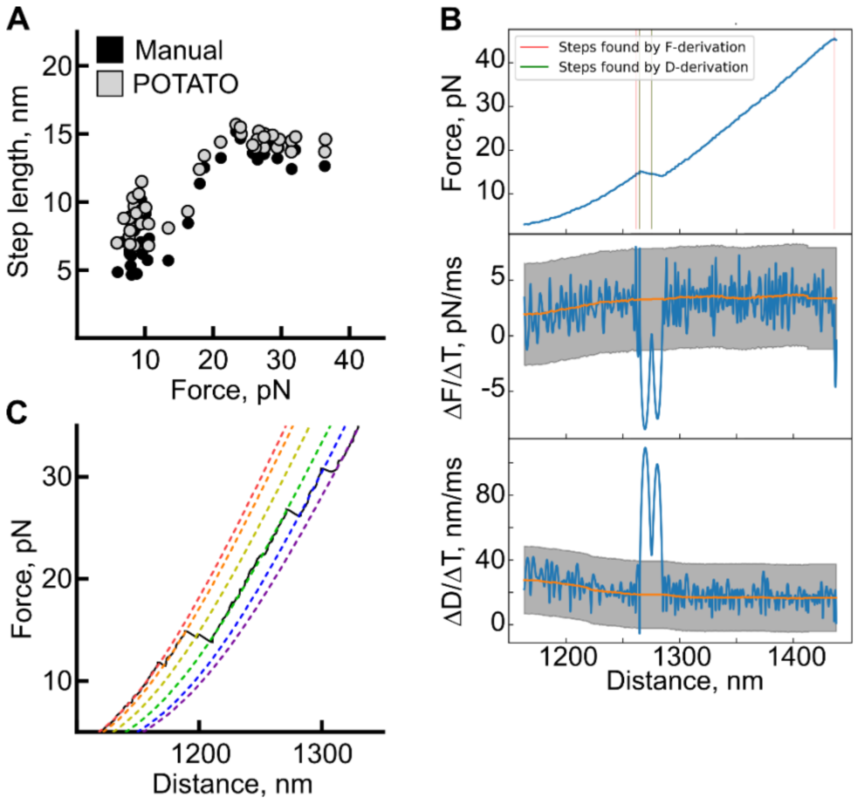


**FIGURE S2: Real data application of POTATO**. **(A)** Comparison of unfolding events marked in a subset of the data analysed manually (black) or with POTATO (grey). . **(B)** Example analysis output from POTATO showing the trimmed FD curve (up), force derivative data (middle), and distance derivative data (bottom). An intermediate conformer is detected by POTATO during the unfolding. Other FD curves confirmed the presence of an even more stable and distinct intermediate step. (**C**) Example curve (black, solid) with five unfolding steps fitted by POTATO (colored, dashed).

**TABLE S1: Parameters used throughout the pipeline and a short description.**

| Parameter | Description |
|---|---|
| **Preprocessing** | |
| Downsampling rate | Only every nth value is taken for analysis; speeds up subsequent processing. |
| Butterworth filter degree | Defines the stringency of the filter. |
| Cut-off frequency | Signals with a frequency above this threshold are suppressed. |
| Force threshold, pN | Values with lower force are excluded from the analysis. |
| **Derivative** | |
| Step d | Characterizes the interval between two values used for numerical derivative calculation. |
| Data frequency, Hz | Frequency at which data is recorded. |
| **Statistics** | |
| Z-score | The number of standard deviations used to determine whether a given value is part of a normal distribution. |
| Moving median window size | The number of values considered for each median calculation. |
| SD difference threshold | Statistical analysis and data sorting are iterated until the difference between two consecutive SDs is below this value. |
| **Fitting** | |
| dsLp, nm | Persistence length of the double-stranded (folded) part of the tethered construct. |
| dsLc, nm | Contour length of double-stranded (folded) part of the tethered construct. |
| dsK0, pN | Stretch modulus of double-stranded (folded) part of the tethered construct. |
| Force offset, pN | Force offset of a given dataset; compensates for a shift in the dataset. |
| Distance offset, nm | Distance offset of a given dataset; compensates for a shift in the dataset. |
| ssLp, nm | Persistence length of the single-stranded (unfolded) part of the tethered construct. |
| ssLc, nm | Contour length of single-stranded (unfolded) part of the tethered construct. |
| ssK0, pN | Stretch modulus of single-stranded (unfolded) part of the tethered construct. |

**TABLE S2: Dependence of the performance measures on the z-score**. Analysis of 2520 generated curves with steps occurring between 10-40 pN with different z-score values.

| z-score / Parameter | 3.2 | 3 | 2.7 | 2.5 |
|---|---|---|---|---|
| True positives | 1206 | 1267 | 1280 | 1303 |
| True negatives | 1101 | 1076 | 1073 | 1056 |
| False positives | 32 | 57 | 60 | 77 |
| False negatives | 181 | 120 | 107 | 84 |
| Accuracy | 0.915 | 0.930 | 0.934 | 0.936 |
| Precision | 0.974 | 0.957 | 0.955 | 0.944 |
| Recall | 0.870 | 0.913 | 0.923 | 0.939 |
| Specificity | 0.972 | 0.950 | 0.947 | 0.932 |
| F1-Score | 0.919 | 0.935 | 0.939 | 0.942 |

**TABLE S3: Python packages used in POTATO.** Standard packages are not included in the table.

| Package name | link |
|---|---|
| h5py | https://www.h5py.org (2) |
| Pandas | https://pandas.pydata.org (3) |
| Scipy | https://www.scipy.org (4) |
| Matplotlib | https://matplotlib.org (5) |
| Lumicks.pylake | https://lumicks-pylake.readthedocs.io |
